# Totem: a user-friendly tool for clustering-based inference of tree-shaped trajectories from single-cell data

**DOI:** 10.1101/2022.09.19.508535

**Authors:** Johannes Smolander, Sini Junttila, Laura L Elo

**Author notes:** To whom correspondence should be addressed **Contact information** Turku Bioscience Centre, University of Turku and Åbo Akademi University, FI-20520 Turku, Finland.

## Abstract

Single-cell RNA-sequencing enables cell-level investigation of cell differentiation, which can be modelled using trajectory inference methods. While tremendous effort has been put into designing these methods, inferring accurate trajectories automatically remains difficult. Therefore, the standard approach involves testing different trajectory inference methods and picking the trajectory giving the most biologically sensible model. As the default parameters are often suboptimal, their tuning requires methodological expertise. We introduce Totem, an open-source, easy-to-use R package designed to facilitate inference of tree-shaped trajectories from single-cell data. Totem generates a large number of clustering results, estimates their topologies as minimum spanning trees, and uses them to measure the connectivity of the cells. Besides automatic selection of an appropriate trajectory, cell connectivity enables to visually pinpoint branching points and milestones relevant to the trajectory. Furthermore, testing different trajectories with Totem is fast, easy, and does not require in-depth methodological knowledge.

## Main

Single-cell RNA-sequencing (scRNA-seq) enables researchers to quantify the transcriptome of thousands or even millions of cells simultaneously at the single-cell level. The cells in a tissue undergo transcriptomic changes that are part of biological processes. If the changes happen gradually, the tissue contains cells derived from different stages of the process. If the correct trajectory that accurately models these different stages can be generated from the scRNA-seq data, this enables researchers to unravel the transcriptomic mechanisms of the processes. The modelling of trajectories from scRNA-seq data has formed a new branch in computational biology, known as trajectory inference.

A vast number of trajectory inference methods have been developed^1,2^. While these methods have proven to be valuable in many situations, their use is still difficult for several reasons. The main reason is that a single method used with default settings often fails to generate a trajectory that can capture all relevant parts (milestones) of the trajectory and also accurately model the correct milestone network. Therefore, a popular approach is to test different methods and tune their parameters until a biologically sensible or otherwise satisfactory trajectory is obtained.

Slingshot^3^ has become one of the most popular trajectory inference methods after a comparison study^2^ showed it was one of the best-performing methods for inferring tree-shaped trajectories. A tree can be any trajectory in which the parts are not disconnected and the trajectory has no cycles. Slingshot requires as input a clustering of cells that is used to construct a Minimum Spanning Tree (MST), which is subsequently smoothed using the simultaneous principal curves algorithm to obtain a directed trajectory, which also includes pseudotime, a measure of the differentiation stage of the cells. While methods such as the Average Silhouette Width (ASW)^4^ and the Variance Ratio Criterion (VRC)^5^ can be used to select the clustering automatically, their weakness is their tendency to select a clustering with a too small number of clusters. The resulting trajectories are often over-simplistic and lack especially small milestones. Therefore, a more reliable approach can be to manually generate a clustering that includes all the relevant milestones, but this requires careful parameter testing and validation using markers to ensure that the clustering is optimal. However, even then the resulting MST may not correlate well with the true milestone network because the MST is generated using a distance matrix that is sensitive to the pre-processing steps (e.g. dimensionality reduction), the clustering structure, and the choice of the distance metric.

To provide a system with better user experience for researchers who perform trajectory inference, we developed a new tool, Totem, for the inference of tree-shaped trajectories from single-cell data (**Fig. 1**). Totem generates a large number of dissimilar clustering results for the cells (by default, 10,000) using a *k*-medoids algorithm. The number of clusters (*k*) and the structure of the clustering results vary, generating vastly different trajectories when used as the basis for constructing the MST. To select a clustering from the large set of clustering results, Totem uses a new measure called the cell connectivity, which is based on counting the number of edges (connections) between the clusters in an MST. The ratio of the number of edges and clusters (connectivity) is calculated for each cell and clustering, and the connectivity vectors are averaged to obtain the cell connectivity of the cells. The cell connectivity acts as a useful baseline for selecting an appropriate trajectory and deciding which milestones to include in the trajectory, while also helping to give a visual overview of the milestone network, i.e. how the milestones are connected and where the branching points and leaf nodes are located. Importantly, a key feature of Totem for its usability is that it allows to quickly and easily browse different MSTs, from which the user can select one or several MSTs for further analysis.

**Fig. 1:**
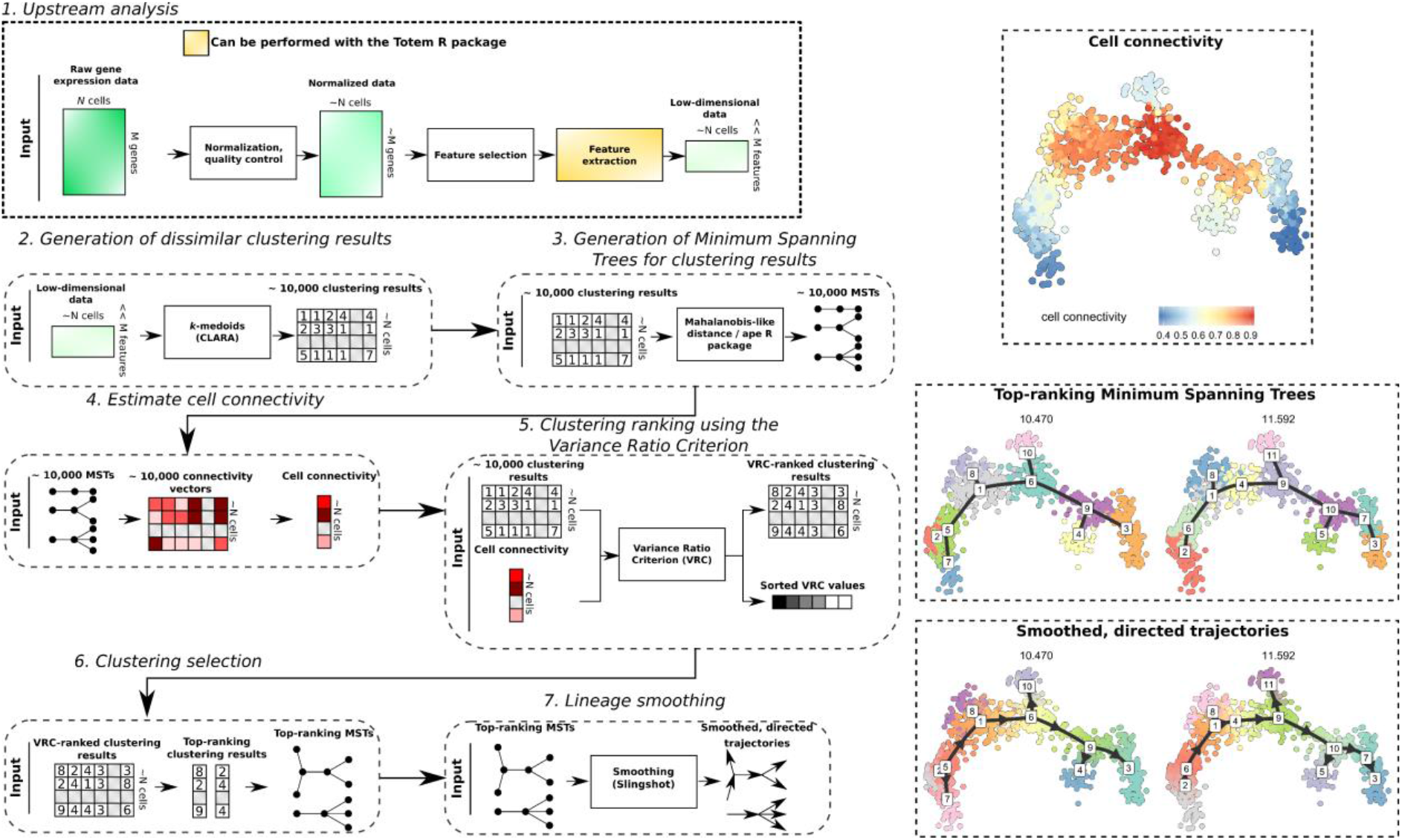
Schematic illustration of the Totem workflow. **(1)** A gene expression matrix is used as input, which needs to be pre-processed upstream of Totem analysis, including normalization, quality control, and possible batch correction and feature selection. Further dimensionality reduction (feature extraction) can also be performed using Totem. For the low-dimensional data matrix, Totem generates **(2)** a large set of clustering results using a *k*-medoids clustering algorithm (CLARA), **(3)** a Minimum Spanning Tree (MST) for each clustering, which models the cluster (milestone) network, and **(4)** estimates the connectivity of the clusters in each MST (ratio of the number of edges a cluster has in MST and the number of clusters). The connectivity vectors are averaged to generate the cell connectivity measure (top right), which helps to locate branching points and milestones that are relevant to model. **(5)** The clustering results are ranked based on the Variance Ratio Criterion (VRC) of the cell connectivity measure, and **(6)** top-ranking clustering results and their corresponding MSTs are selected for further analysis (middle-right). **(7)** The selected MSTs are smoothed using the Slingshot algorithm to obtain directed trajectories (lower right). The trajectory can be exported as objects that can be used downstream of Totem analysis, such as differential expression analysis.

To benchmark Totem, we performed a comprehensive comparison using the dynverse benchmarking framework^2^, which comprises over 200 tree-shaped trajectories. We compared Totem with the popular Slingshot tool^2,3^ and the more recently introduced TinGa^6^, which is based on a growing neural gas (GNG) model^7^ and can also model trajectories that are more complex than trees. In particular, to evaluate the performance of the connectivity-based criterion of Totem for clustering selection in trajectory inference, we compared it with the popular ASW and VRC clustering selection methods.

Because a single clustering is rarely optimal, and thus the users are often forced to try multiple clustering results to optimize the trajectories, we investigated the performance of the trajectories generated based on multiple top-ranked clustering results, giving a deeper overview of the performance. Finally, we demonstrate the usefulness of the cell connectivity measure with a few examples that show how the measure can aid trajectory inference by helping to pinpoint relevant branching points and milestones.

## Results

### Benchmarking Totem for trajectory inference

To benchmark Totem, we used the dynverse benchmarking framework^2^ that includes 216 tree-shaped trajectories. In our comparison, we included the dynverse implementation of Slingshot^2,3^, which uses the ASW method for clustering selection, and TinGa, which is a GNG-based method^7^ that can also predict more complex trajectories than tree-shaped, such as acyclic and disconnected trajectories^6^.

As shown in the rightmost panel of **Figure 2**, Totem achieved superior overall scores (Wilcoxon signed-rank test; *P*-value ≤0.01) for datasets with a non-linear trajectory (bifurcating, multifurcating, or other non-linear tree), whereas TinGa was the second-best method for the non-linear datasets. The violin plots also show that Totem’s distribution of the overall scores with the non-linear trajectories was more concentrated to higher performance levels compared to TinGa and Slingshot. For linear trajectories, Slingshot achieved superior performance among the tested methods (Wilcoxon signed-rank test; *P*-value ≤ 0.01), while the overall scores of the linear datasets were, on average, similar for TinGa and Totem. The difference in the overall performance was mainly attributable to the differences in the topology accuracy (HIM) and the accuracy of the cell assignment onto the branches (F1 branches). The HIM score of Slingshot was close to perfect for most of the linear datasets but worse compared to TinGa and Totem for the non-linear datasets.

**Fig. 2.**
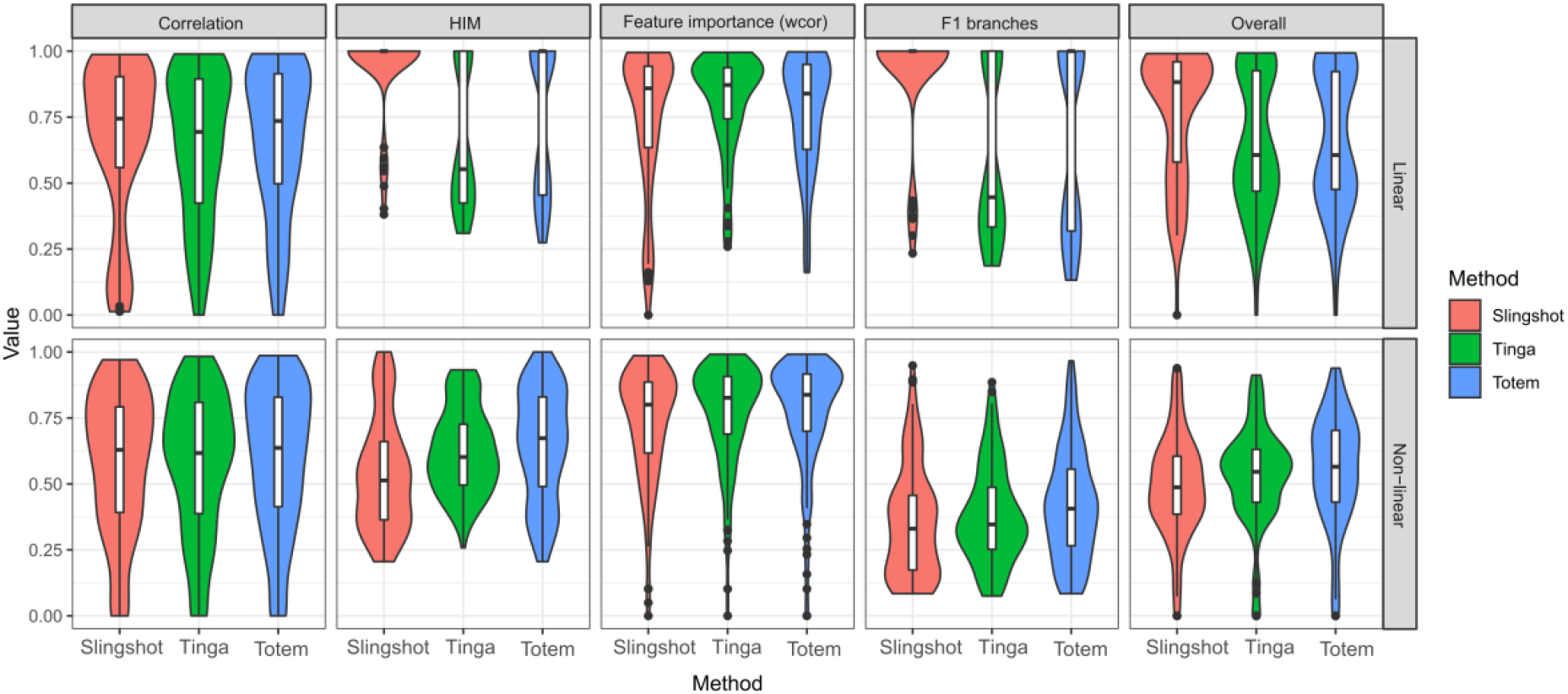
Results of the benchmarking for trajectory inference from scRNA-seq data. The overall score is the geometric mean of four performance metrics: the correlation of geodesic distances (correlation), which measures accuracy of cellular ordering, the Hamming-Ipsen-Mikhailov (HIM), which measures topological accuracy, the weighted correlation of feature importance lists, which measures accuracy of differentially expressed genes inferred from the trajectory, and the F1 branches, which measures accuracy of cell assignment onto branches. The results were grouped by the performance metric (columns) and whether the ground-truth topology is linear or non-linear, i.e. bifurcating, multifurcating, or some other non-linear tree (rows).

### Benchmarking Totem for clustering selection

Selection of an appropriate clustering for constructing the MST is an important step in the trajectory inference process. Therefore, we took a closer look at the clustering selection step to investigate which clustering selection approach gives the best results when we consider multiple top-ranking trajectories simultaneously. We included two popular clustering selection methods, which were the average silhouette width (ASW) and the Variance Ratio Criterion (VRC), as well as the Totem method that uses the cell connectivity and the VRC, and also a method that ranks the clustering results into a random order (Random).

When we tested multiple top-ranking clustering results and always selected the trajectory that gave the best performance in the evaluation, the result suggested that the random criterion gave the best performance (**Fig. 3A**). This is an expected result considering that the random criterion generates more dissimilar trajectories than the other methods. However, when we averaged the performance values of the multiple top trajectories (**Fig. 3A**), the random criterion was the worst-performing method, as expected. The ASW achieved the worst performance among the methods in the analysis in which we selected the best-performing trajectory, and it was the second-worst method when the performance values of the trajectories were averaged. The VRC achieved slightly better performance than the ASW when selecting the best-performing trajectory, and it was tied with the Totem method as the best method by the average performance. In addition to good average performance, Totem had a relatively high performance curve in the analysis that considers the performance of the best-performing trajectories, similar with the random criterion. In other words, the selection method of Totem enables to find better performing trajectories than the ASW and VRC without sacrificing as much on the average performance as the random criterion.

**Fig. 3.**
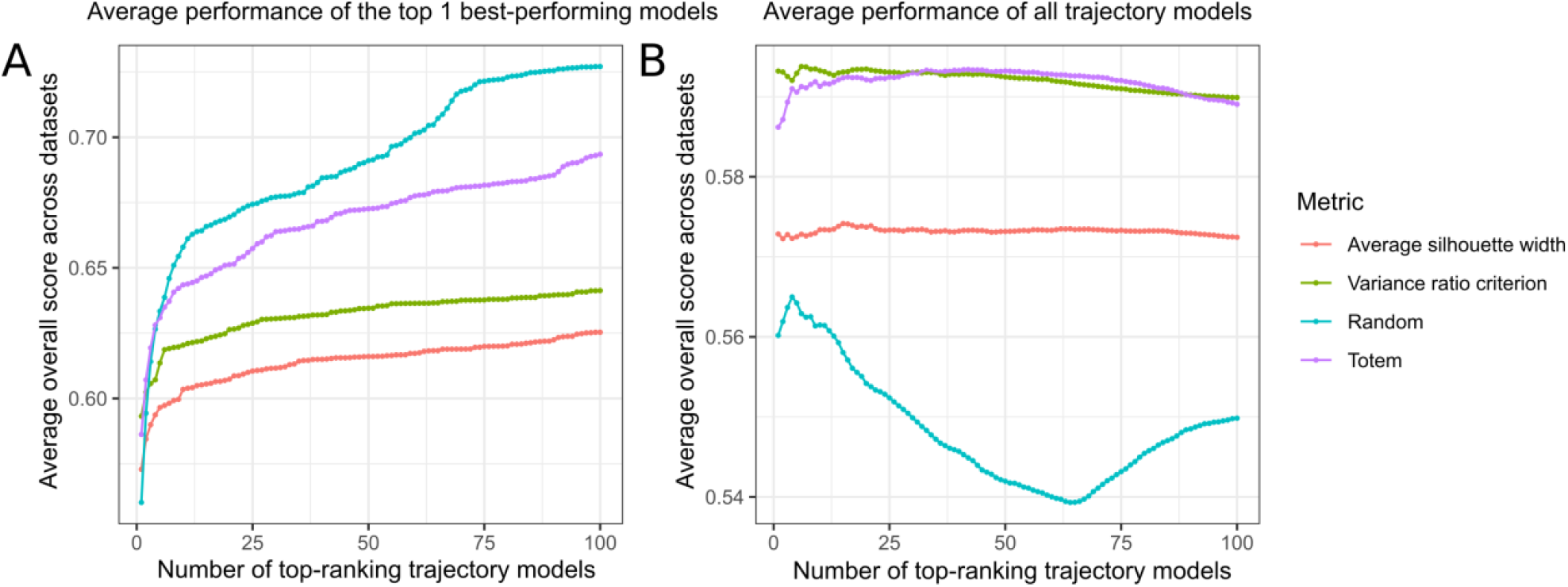
Comparing clustering selection methods in the dynverse benchmarking. We selected the top 100 clusterings from 10,000 k-medoids clusterings for each of the 216 dynverse benchmark datasets using four different methods: The average silhouette width, the Calinski-Harabasz score, random selection, and the Totem method. For each clustering, the rest of the trajectory inference (MST generation, smoothing) was performed using Slingshot. In (A), we varied the number of top- ranking trajectory models and selected the model for each dataset that gave the best overall score from all the models. In (B), we calculated the mean performance of all the models.

### Examples of trajectory inference with Totem

To demonstrate the usefulness of Totem in practice, we compared the trajectories (**Fig. 4A**) produced by Totem and Slingshot for a simulated dataset that has a multifurcating trajectory (named multifurcating_4 in the dynverse benchmark data repository). With Slingshot, we used the ground-truth clustering of the simulated dataset as input. The MST produced by Slingshot was linear and did not correlate well with the true milestone network. The example shows how Slingshot can still produce inaccurate trajectories even if the ground-truth clustering is available because the cluster distances generate an MST with a wrong network. In contrast, Totem predicted trajectories that correlated well with the true milestone network. In addition, the cell connectivity measure of Totem provided, in general, accurate information about the location of the end points and the middle branching point.

**Fig. 4.**
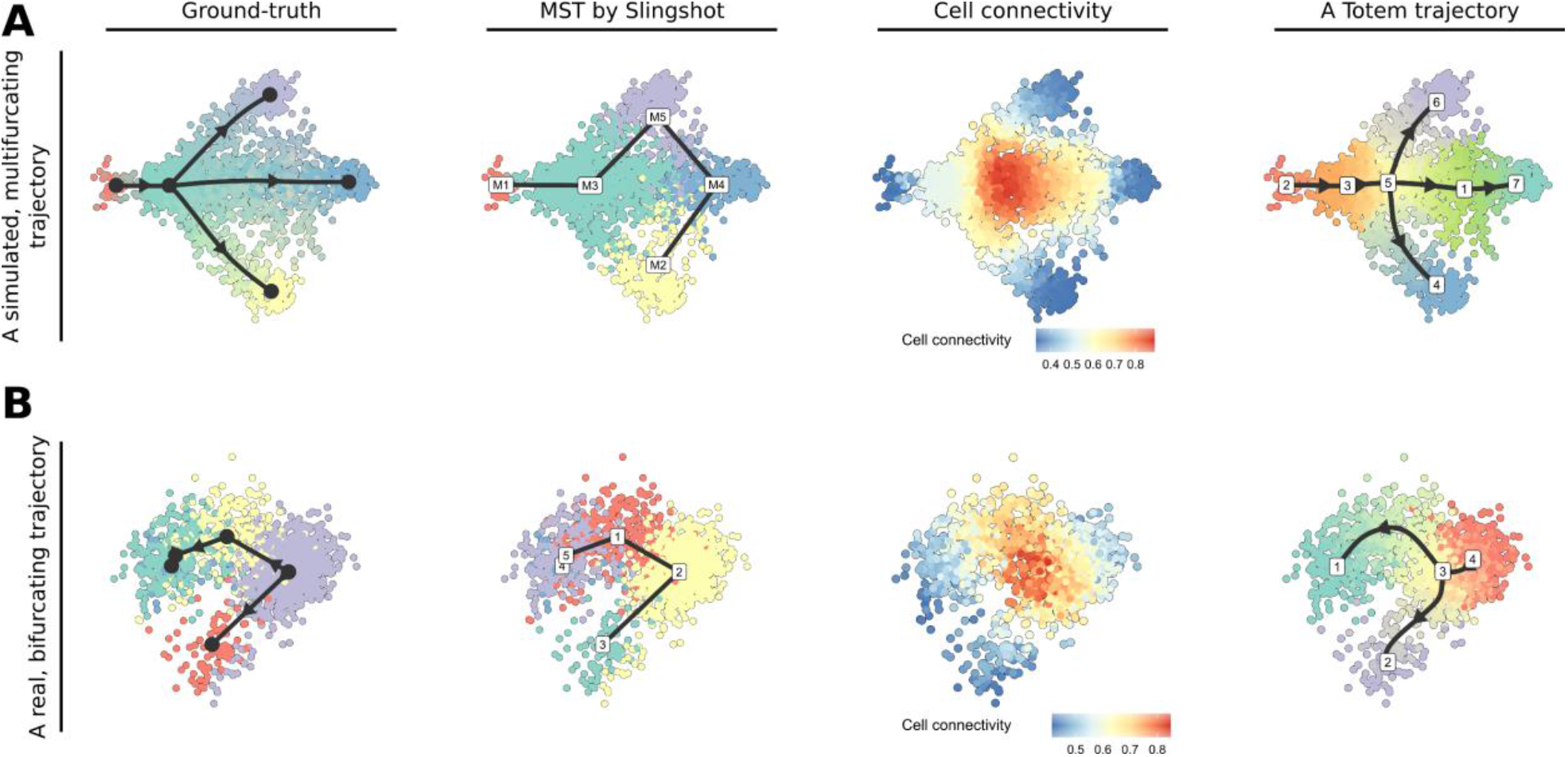
Example analyses with Slingshot and Totem. **(A)** A multifurcating trajectory simulated with dyntoy. **(B)** T cells from a mouse thymus with a bifurcating trajectory. The two-dimensional embeddings for visualization were generated with the Multi-Dimensional Scaling (MDS) method from the dyndimred R package.

The second example (**Fig. 4B**) shows similar results but for a bifurcating trajectory that includes T cells from a mouse thymus^2,8^. Based on the trajectory inferred by Slingshot, the topology of this network could be linear, i.e. the start point would be one of the end nodes, or bifurcating, i.e. the start point would be one of the middle nodes, if we have no knowledge about the starting point of the trajectory. By selecting a trajectory with Totem that is in line with the cell connectivity profile of the data, we obtain a trajectory with the correct topology (bifurcating), where the bifurcation point is at the node in the middle, from which the bifurcation starts in the ground-truth trajectory. Both of these examples demonstrate, how the cell connectivity helped us to choose trajectories that exhibit the correct topology.

## Discussion

scRNA-seq can be a powerful technology for studying transcriptomic processes of cellular systems. While a large number of trajectory inference methods has been developed^2^ for constructing trajectories from single-cell data through mathematical modelling, the usability of these methods remains rather poor. One of the main reasons is that the methods do not work optimally with every dataset, requiring the users to try different methods, tune the parameters, and adjust the pre-processing steps prior to trajectory inference. While helpful frameworks like dyno^2^ have been developed that enable testing different trajectory methods, it still takes considerable effort to fine-tune the trajectory inference models. Oftentimes, the parameter tuning requires detailed knowledge of the underlying statistical and machine learning models, which many users are not familiar with, and the manuals of the trajectory inference tools often do not provide clear instructions on how the parameters should be adjusted.

In this paper, we introduced Totem, a tool designed to facilitate the inference of tree-shaped trajectories from single-cell data. Totem generates a large number of clustering results (by default, 10,000) with the *k*-medoids algorithm and calculates the connectivity of the clusters, i.e. the ratio of the connected clusters and the total number of clusters, in the minimum spanning trees (MSTs) that are generated from the clustering results. By averaging the connectivity vectors of all the clustering results, we obtain the cell connectivity measure, which acts as a useful baseline for finding the optimal trajectory. The cell connectivity can be visualized over a two-dimensional embedding, such as the multi-dimensional scaling (MDS) or *t*-distributed stochastic neighbour embedding (*t*-SNE), and the local changes in the connectivity indicate milestones and branching points that are relevant to include in the trajectory model. In our examples, we showed how the cell connectivity enabled us to select clustering results that generated accurate trajectories.

Other trajectory inference methods, such as Slingshot^3^ and TinGa^6^, do not provide a metric like the cell connectivity that can be used as a reference to aid trajectory inference. With these methods the user needs to instead use the visualization to get a sense of the topology, use biological markers to ensure the model is biologically sensible, and change the parameters if the trajectory is not satisfactory, which can be arduous for users without advanced background in mathematics and machine learning.

In Slingshot, the between-cluster distances determine the MST and the milestone network. Our examples showed how even when the correct cell types were available, the MSTs predicted by Slingshot could still be inaccurate. One solution is to change the distance method, for example, to a mutual neighbor -based method, but this will still not necessarily generate the correct network. Totem addresses this issue by providing a user-friendly interface to analyze MSTs generated based on different clustering results. With this interface the user can select an MST that best fits to the biological hypothesis and the cell connectivity profile without time-consuming and arbitrary parameter tuning. Totem then smoothens the MSTs of the selected clustering results using the simultaneous principal curves algorithm of Slingshot to obtain directed trajectories that include the pseudotime. The trajectories can be exported as Slingshot or dynwrap objects to be used in downstream analysis, for example to perform differential expression analysis using tradeSeq^9^ or dyno^2^.

To benchmark Totem for inferring trajectories from scRNA-seq data, we used the dynverse benchmarking framework, which includes 216 tree-shaped trajectories. We compared Totem with the dynverse implementation of Slingshot, which uses *k*-medoids to cluster cells and the average silhouette width (ASW) to select the optimal clustering, and TinGa, which is a growing neural gas (GNG) -based method. For clustering selection with Totem, we used the combination of the cell connectivity measure and the variance ratio criterion (VRC). Totem outperformed the other two methods for non-linear trajectories (bifurcation, multifurcation, or some other non-linear tree) and performed comparably with TinGa for linear trajectories. The results suggested that while Slingshot was the best method for linear trajectories among the tested methods, it was significantly worse than TinGa and Totem for non-linear trajectories. This happened because the ASW is generally known to select a small number of clusters, which will more likely generate a linear trajectory than a non-linear trajectory when used for MST estimation. By contrast, TinGa is more likely to overcomplicate the trajectories with redundant cycles and disconnected parts than Slingshot and Totem, both of which can only infer tree-shaped trajectories. Indeed, the performance differences between the methods were mainly attributable to differences in the accuracy of the topology (HIM) and branch assignment (F1 branches).

To investigate what is the most optimal way to select the clustering that is used to generate the MST, we generated 10,000 *k*-medoids clustering results for each of the benchmark datasets and ranked them using different clustering evaluation methods, including the ASW, the VRC, and the Totem criterion that uses the cell connectivity and the VRC. When we investigated the performance scores of the top 100 clustering results, the results suggested that the Totem criterion and the VRC provided the best average performance. However, the Totem criterion outperformed the VRC when we considered only the best trajectory from the set of trajectories. In other words, when testing multiple trajectories, it is more likely that we find a higher-performing trajectory with Totem than with the ASW and VRC methods, and the average performance of all the trajectories found by Totem will likely be at least as good as for the other methods.

Totem has the same limitations as Slingshot. In particular, it cannot be used to infer trajectories that have cycles or both diverging (cells diverge from a single point into several lineages) and converging (cells converge into a single point from several lineages) parts at the same time. In addition, unlike the updated Slingshot, Totem cannot currently handle disconnected trajectories. However, we do not consider this a major limitation because determining automatically which cell types belong to which disconnected sub-trajectories is not an easy task^10^. A safer approach is to analyse the disconnected parts as separate trajectories by segregating the cell types manually before trajectory inference.

Although there exist methods that can handle almost arbitrary topologies, such as PAGA^11^, Monocle3^12^, and TinGa^6^, their issue is that they are more likely to overcomplicate the trajectory with extra cycles and branches than methods like Slingshot and Totem that are limited to tree-shaped trajectories. Similarly, methods like scShaper^12^, SCORPIUS^13^, and Elpilinear^14^ that are limited to linear trajectories are still useful because they are guaranteed to provide the correct topology if the trajectory is expected to be linear, unlike the more complex methods.

To summarize, Totem is a tool designed to facilitate the inference of tree-shaped trajectories from single-cell data. It is built upon the popular Slingshot method, which uses a clustering to construct an MST and the simultaneous principal curves algorithm to obtain a directed trajectory along with pseudotime that quantifies cell differentiation at the single-cell level. The benefit of Totem over the available tools is that it is designed to provide a user-friendly interface for finding a clustering that is used as the basis of the trajectory. Although the clustering selection is highly critical for the success of the trajectory inference, performing it automatically in a way that will generate the correct milestone network in the trajectory remains challenging. To address this challenge, the analysis of different clustering results and their MSTs with Totem has been designed to be fast and easy without requiring complex parameter tuning and in-depth technical knowledge. As a notable difference compared to existing trajectory inference methods, Totem provides the cell connectivity measure, which aids the trajectory optimization by providing information about the location of the branching points and milestone transitions.

## Methods

### Totem

In the following, we go through the main steps of the Totem trajectory inference workflow. **Figure 1** illustrates the basic workflow of Totem.

### Upstream analysis

Upstream of trajectory inference, Totem assumes that the input gene expression matrix has been normalized and quality control has been performed to remove bad-quality cells^15^. scRNA-seq data analysis toolkits like Seurat^16^ and Scanpy ^17^ can perform the normalization and quality control.

Trajectory inference methods require a low-dimensional (from 2 to 50) embedding as input. The dimensionality reduction steps can be customized as the user sees best prior to Totem analysis, but Totem can also be used to transform gene counts into new features (feature extraction) with functions that utilize the dyndimred R package. In scRNA-seq data analysis, the most common way to perform feature extraction is the Principal Component Analysis (PCA) in which the number of principal components typically varies from 10 to 30. Alternatively, methods like Multi-Dimensional Scaling (MDS) or the more scalable landscape MDS (LMDS) can be used to generate an embedding with a smaller number of dimensions (e.g. from 2 to 5), which is usually not a sufficient number in PCA of scRNA-seq data. *t*-SNE^18^ and UMAP^19^ are commonly used only for visualization, and they should be used with caution if used as input in trajectory inference. As the default dimensionality reduction method in Totem, we use the 5-dimensional LMDS. Moreover, it is generally a good idea to reduce the number of features prior to feature extraction by selecting highly variable genes (HVGs), which can be performed using methods like Seurat ^16^ and Scanpy^17^. If the dataset has batch effects, and the user does not want to model these differences in trajectory inference, data integration methods ^16,20^ can be used.

### Generation of a large set of clustering results

To generate a large set of dissimilar clustering results that can be used as the basis for constructing the Minimum Spanning Tree (MST), we use the CLARA *k*-medoids clustering algorithm^21^ from the cluster R package^22^, which performs fast *k*-medoids clustering. We run the algorithm *L* times (by default *L* = 10,000) and filter out clustering results that have clusters with fewer than 5 cells to prevent overly small clusters, which are usually not interesting. The number of remaining clustering results (*L*′) can be close to *L* or deviate from it considerably depending on the dataset. The number of clusters (*k*) varies by default from 3 to 20, but can be adjusted by the user if more complex trajectories with more milestones are expected. The default upper limit (20) is well above the average number of clusters (milestones) in the trajectories of the dynverse benchmarking framework. We also activate the R random number generation (RNG) in the clara R function so that the program returns different results with different RNG seeds.

### Generation of Minimum Spanning Trees for clustering results

For each clustering, we generate an MST by representing the clusters of cells as vertices in a graph and using the covariance-based approach of Slingshot^3^ to measure distances between the clusters. The Mahalanobis-like distances are calculated from the covariance matrix for cells in each cluster, while also considering the shape and spread of the clusters. We use the ape R package to perform the MST estimation23.

### Estimate cell connectivity

For each MST *j* (*j* = 1, …, *L*′), we measure the connectivity *cij* = *dij/kj* of the ith vertex (cluster) in the graph by dividing the degree of the vertex *dij*, i.e. the number of edges that are connected to the vertex, by the number of vertices *kj* in the graph. We then create a connectivity vector ***c****j* that contains the connectivity values for all *N* cells corresponding to MST *j*, scale the connectivity values so that the maximum connectivity of each graph is always one, and calculate the cell connectivity of all the cells 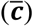 by averaging over the *L*′ connectivity vectors:

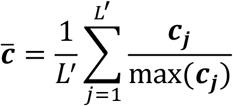

The cell connectivity is higher for cell populations that are farther from the leaf parts of the trajectory (**Fig. 1**).

### Clustering selection using the variance ratio criterion

To find a clustering that captures the tree structure of the data and gives clusters that are well defined, we use the variance ratio criterion (VRC), also known as the Calinski-Harabasz score^5^, which measures the ratio of the between-cluster dispersion and the within-cluster dispersion. Instead of using the whole dimensionally reduced data as input, we use the one-dimensional cell connectivity vector 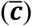 to find clusters that have cells with similar connectivity within the clusters but different compared to the other clusters. The VRC is a more memory-efficient method for clustering selection than the commonly used average silhouette width (ASW), because it does not require a distance matrix of cells as input, making it more suitable for datasets with many cells. We use the fpc R package to perform the VRC analysis^24^.

### Lineage smoothing

For an MST corresponding to a selected clustering, Totem performs lineage smoothing using Slingshot^3^, which generates a directed trajectory along with pseudotime that quantifies cell differentiation continuously at the single-cell level. Slingshot fits principal curves^25^ for each lineage of the MST starting from a user-specified root node, which is initially selected randomly. The root node can be adjusted later by the user.

### Trajectory visualization and interpretation

Totem includes visualization functions that aid trajectory inference, utilizing the dynplot R package^2^. The functions enable to visualize multiple MSTs and smoothed trajectories side by side over a two-dimensional embedding. The two-dimensional embedding can be provided by the user or generated using one of the methods included in the dyndimred R package, such as *t*-SNE, UMAP, or MDS. To assist in biological interpretation, the user can also visualize expression levels of genes over the embedding.

The cell connectivity can be visualized over the two-dimensional embedding to pinpoint milestones and branching points that could be otherwise missed or inaccurately modelled **(Fig. 1)**. Local changes in the connectivity indicate milestone transitions that are relevant to include in the trajectory. The connectivity levels can be used to locate branching points, where a cell population is encircled by other cell populations whose connectivity is relatively lower **(Fig. 4)**.

Although the clustering selection is performed based on the VRC of the cell connectivity, the resulting MST can still be suboptimal in terms of the connections that the milestone network comprises. In addition, small milestones can be missing or the trajectory can seem over-complicated. Therefore, the user should not rely upon a single, automatically generated trajectory. Instead, the trajectory should be validated using gene markers to ensure that the model is biologically sensible, and compared with the cell connectivity by visualization to ensure that the milestone network is in line with the connectivity profile. If the trajectory requires adjustments, Totem allows to easily test different clustering results until a trajectory is obtained that meets both requirements.

### Benchmarking

To benchmark Totem, we repeated the comprehensive benchmarking of scRNA-seq trajectory inference methods using the dynverse framework2. We included trajectories that have a tree-like structure, which were labeled as linear, bifurcation, multifurcation, or tree in the original comparison. These comprise 69, 44, 16, and 87 linear, bifurcation, multifurcation, and tree trajectories, respectively. The datasets are of synthetic and real origin. The real datasets include 26 gold standard datasets of which ground-truth includes cell types and their differentiation order at the cluster level (discrete pseudotime). The 54 real, silver standard datasets have a ground-truth that includes a continuous or discrete pseudotime. The synthetic, simulated datasets provide the most accurate ground-truth, and they were simulated using four different simulators: dyntoy^2^, dyngen^26^, PROSSTT^27^, and Splatter^28^.

The dynverse benchmarking framework includes four main metrics. The accuracy of the cell differentiation order is measured using the correlation of pairwise geodesic distances between the ground-truth and predicted trajectories. The accuracy of the differentially expressed genes is measured using the weighted correlation of the random forest regression -derived feature importance lists. The features that are ranked higher in the ground-truth are given a relatively higher weight. The F1 branches value maps the cells to the closest branching points in the ground-truth and inferred trajectories and measures the clustering similarity between the two clustering results. Hamming-Ipsen-Mikhailov (HIM)^29^ measures the similarity of two topologies. For example, two linear trajectories would yield a perfect HIM value of 1, even if the two trajectories would be otherwise completely dissimilar. All four metrics range from 0 to 1 and the geometric mean of the metrics is used to assess the overall performance by penalizing small values in the metrics.

To benchmark Totem, we compared it with the dynverse implementation of the Slingshot method, which uses the PAM (*k*-medoids) clustering algorithm and the average silhouette width to select the optimal number of clusters, as well as TinGa, which is based on a growing neural gas model. However, the pre-processing was standardized so that each method uses the 5-dimensional latent multidimensional scaling (LMDS).

### Comparing clustering selection methods

Trajectory inference methods such as Slingshot require a clustering that is used as the basis for constructing the MST, which is subsequently smoothed using the simultaneous principal curves algorithm to obtain a directed trajectory and pseudotime. In addition to the cell connectivity criterion of Totem for clustering selection, we tested three other methods: the average silhouette width (ASW), which is used by the dynverse implementation of Slingshot; the Calinski-Harabasz score, also known as the Variance Ratio Criterion (VRC); and the random criterion that ranks the clustering results into a random order.

To compare the clustering selection methods, we ran the *k*-medoids clustering algorithm (CLARA) 10,000 times for each dataset, selected the top 100 clustering results with each method, performed the trajectory inference using Slingshot for each clustering, and ran the dynverse performance evaluation for the trajectories. For each benchmark dataset, we varied the number of evaluated trajectories from 1 to 100 and calculated the maximum and mean of the overall performance score. Finally, we calculated the mean of the overall scores across the datasets.

## Data availability

The benchmark data is available on Zenodo30.

## Code availability

The Totem R package is available at https://github.com/elolab/Totem, and it includes a vignette that explains how Totem should be used. The codes that are relevant for repeating the benchmarking are available at https://github.com/elolab/Totem-benchmarking.

## Acknowledgements

The authors thank the Elo lab for fruitful discussions and comments on the manuscript.

Prof. Elo reports grants from the European Research Council ERC (677943), European Union’s Horizon 2020 research and innovation programme (955321), Academy of Finland (310561, 314443, 329278, 335434, 335611 and 341342), and Sigrid Juselius Foundation, during the conduct of the study. Our research is also supported by University of Turku, Åbo Akademi University, Turku Graduate School (UTUGS), Biocenter Finland, and ELIXIR Finland.

## Ethics declarations

## Competing interests

The authors declare no competing interests.

## Notes

### Competing Interest Statement

The authors have declared no competing interest.

